# Epigenome-wide Association Study of Psilocybin-Induced Methylome Changes in Alcohol Use Disorder

**DOI:** 10.1101/2025.07.18.664368

**Authors:** Marvin M. Urban, Lea Zillich, Nathalie M. Rieser, Marcus Herdener, Franz X. Vollenweider, Rainer Spanagel, Katrin H. Preller, Marcus W. Meinhardt

## Abstract

The serotonergic hallucinogen psilocybin has shown potential as a treatment for psychiatric conditions like alcohol use disorder (AUD) and depression in clinical studies. Epigenetic mechanisms, including DNA methylation, are hypothesized to contribute to its lasting therapeutic benefits. In this exploratory study, we present the first methylome-wide analysis of psilocybin-induced changes in a cohort of detoxified patients with AUD. The longitudinal study design included three assessment days in 40 patients with blood sampling and acquisition of psychometrics – at baseline, 24 hours after administration of psilocybin (25 mg) or placebo (mannitol), and one month after treatment. Our epigenome-wide association study (EWAS) identified one CpG site in *TLE4* (*p* = 1.1e-7) associated with psilocybin treatment. Screening for differentially methylated regions, we observed altered methylation in the gene *RASGRP4* (*pFDR* = 3.2e-4). Network analysis revealed co-methylation modules related to psilocybin treatment, as well as modules associated with the reduction of depressive symptoms and drinking behavior. Gene ontology analysis indicated involvement of these modules in neuroplasticity and immune functions, suggesting that they may reflect abstinence-related recovery processes. Investigating candidate genes at nominal significance (*p* < 0.05) uncovered promoter-associated methylation changes in *HTR2A* and *TNF*. Furthermore, at *p* < 0.05, we found baseline differences between treatment responders (< 1 standard unit alcohol in 4-week follow-up) and non-responders in genes related to synaptic plasticity and different neurotransmitter systems. While these findings are limited by the modest sample size, they align well with previous literature and might provide starting points for further, large-scale investigations or hypothesis-driven experiments.

## Introduction

Alcohol use disorder (AUD) contributes significantly to the global disease burden, with more than 5% of annual deaths being attributable to excessive alcohol use^1^. Current approved pharmacotherapies for AUD, like acamprosate, disulfiram, or naltrexone, show modest treatment success and require regular dosing, which impedes adherence to therapeutic protocols^2–4^.

Contrasting the dosing regimen of established medications for AUD, psychedelic compounds, such as psilocybin, may be able to induce lasting reductions in alcohol use after a single or few administrations^5–7^. While the underlying mechanisms are not fully understood, recent neurobiological findings suggest a cascade of biological effects involving alterations in gene expression, induction of neuronal plasticity, and changes in functional network connectivity that facilitate shifts in cognition and behavior^8–12^. This chain of effects might also include the level of epigenetics^13,14^.

Several studies support the idea of epigenetic changes in psychedelic drug action. Evidence from preclinical experiments points towards changes in histone acetylation after administration of lysergic acid diethylamide (LSD)^15^ and 2,5-dimethoxy-4-iodoamphetamine (DOI)^16^. Furthermore, a genome-wide methylation analysis after repeated LSD administration revealed 635 differentially methylated cytosine-guanine dinucleotides (CpG sites) in the prefrontal cortex of mice^17^. In a naturalistic human study on ayahuasca, which contains the psychedelic dimethyltryptamine (DMT), increased methylation across five CpG sites in the promoter region of the sigma-1 receptor was found^18^. Apart from that, blood DNA methylation may serve as a marker to predict treatment responses to pharmacological treatments. For instance, methylation patterns measured before pharmacotherapy with antidepressants were associated with treatment success^19,20^.

Here we present the first epigenome-wide analysis study (EWAS) of psilocybin in patients with AUD. Specifically, we investigated longitudinal changes in blood DNA methylation in response to psilocybin treatment in a sample of detoxified patients. The methylation and psychometric data presented here stem from a randomized clinical trial (RCT) that investigated the effect of psilocybin on alcohol relapse and abstinence. This RCT was conducted at the Psychiatric University Hospital in Zurich, Switzerland^21^. While the primary outcomes at 4-week follow-up did not differ between the placebo and psilocybin group – duration of abstinence and mean alcohol use – secondary clinical endpoints, such as depressive symptoms and quality of life, improved significantly in the psilocybin group. Therefore, in the here presented exploratory analysis, we hypothesized i) associations between psilocybin treatment and methylation changes, ii) a mediating effect of methylation changes on depressive symptom reduction in the AUD cohort, and iii) differences in methylation patterns between responders (abstinent at 4-week follow-up) and non-responders to psilocybin treatment. We tested these hypotheses in a methylome-wide manner and for a selection of candidate genes.

## Materials and Methods

### Study Design, Participants, & Psychometrics

This research is based on the study Clinical and Mechanistic Effects of Psilocybin in Alcohol Addicted Patients (clinicaltrials.gov identifier: NCT04141501; Kofam identifier: SNCTP000003445) conducted at the Psychiatric University Hospital in Zürich, Switzerland, by Rieser *et al.*, 2025^21^. The study was a randomized, placebo-controlled, double-blinded, parallel-groups trial, and the final study sample in our analysis included 40 detoxified AUD patients (15 female, 25 male) between 21 to 58 years. The timeline of the trial (Fig. 1) included the acquisition of three blood samples: T1 (baseline, two weeks before dosing visit), T2 (one day after dosing visit), and T3 (around four weeks after dosing visit). During the dosing visit, the treatment group (*n* = 20) was administered 25 mg of psilocybin (acquired from Usona Institute, Madison, Wisconsin) orally while the placebo group (*n* = 20) received an inactive placebo (mannitol).

**Figure 1:**
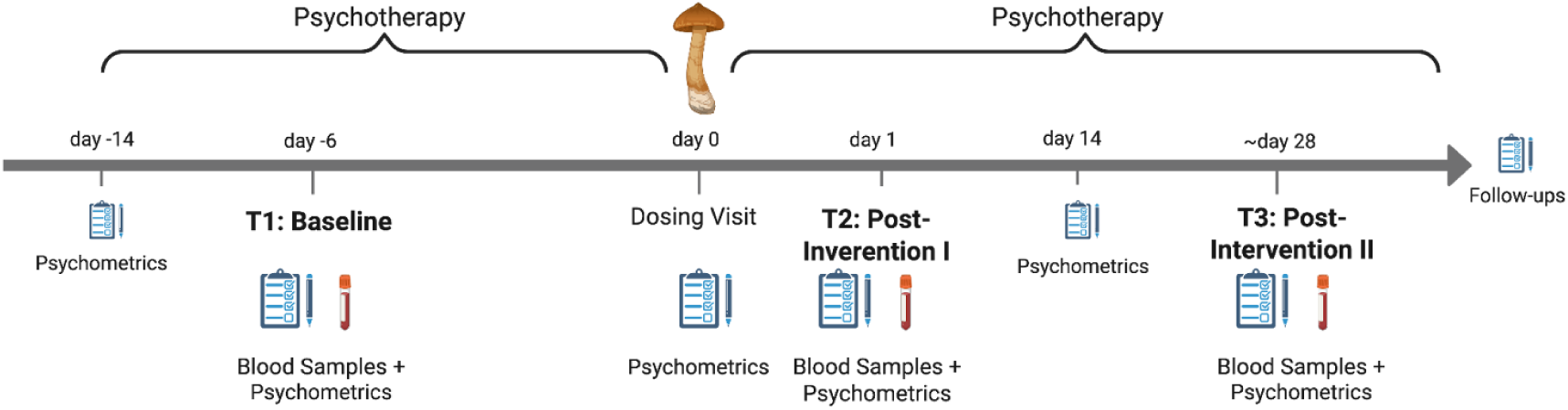
Overview of the clinical trial timeline (*n* = 40), conducted over six weeks. The study included three assessment days with blood sampling and acquisition of psychometrics: baseline (T1), 24 hours post-intervention (T2), and ~28 days post-intervention (T3). Psilocybin (25 mg) or placebo (mannitol) was administered on day 0, the dosing visit, embedded within a broader psychotherapeutic framework incorporating preparatory and integrative sessions. Follow-up assessments were conducted after T3 to monitor longer-term outcomes.

Primary outcomes of interest were daily mean alcohol use in the four weeks after the dosing visit, as well as the time to relapse (≥1 standard unit of alcohol per day). Besides the primary outcomes, we included two secondary psychometric scores: Beck’s Depression Inventory (BDI)^22^ and Beck Hopelessness Scale (BHS)^23^. For these variables, we calculated subject-wise Δ-values between T3 and T1 and used them for downstream analyses of the methylation data. More detailed information on study design, participants, and psychometrics can be found in the original publication^21^.

### Ethical Approval

The clinical part of this study was approved by Swiss legal agencies (Cantonal Ethics Committee, Swiss Agency for Therapeutic Products [Swissmedic], Federal Office of Public Health [BAG]), and adhered to the revised declaration of Helsinki from 2000, as well as guidelines for Good Clinical Practice (GCP). Data sharing and processing adhered to the data protection laws outlined in the General Data Protection Regulation (GDPR) of the European Union.

### DNA Extraction and DNA Methylation Assessment

DNA extraction and methylation analysis were carried out at Life&Brain GmbH in Bonn. 10 ml EDTA-treated whole blood samples were used for DNA extraction via Chemagen Chemagic Systems technology. Extracted DNA was screened for genotypic variants on Illumina’s Infinium Global Screening Array-24 (GSA) v3.0 to include single-nucleotide polymorphism (SNP) outlier analysis in the preprocessing of methylation data. After bisulfite conversion of the DNA, CpG methylation was assessed on Illumina’s Infinium MethylationEPIC BeadChip v2.0, yielding raw data with unmethylated and methylated signal intensities for each of the ~ 950,000 probes stored in idat files, which were then processed as described below.

### Statistical Analysis

All statistical analyses were performed in the *R* statistical environment (version 4.2.1 and 4.3.0; https://www.r-project.org/). A more detailed description of preprocessing and statistical methods can be found in the supplementary materials.

#### Data Preprocessing

Preprocessing was based on the CPACOR pipeline^24^ and included filtering for sample call rate, sex mismatches, genetic outliers, and cross-reactive probes, as described previously^25^. For estimation of genetic outliers, genotype data were preprocessed as previously described^26^, reduced to 20 dimensions by principal component analysis (PCA), and samples for which a component’s loading coefficient differed by 4.5 standard deviations from the mean would have been removed. No such outliers were found. We estimated cell type composition^27,28^ and performed PCA on cell type data and internal control probes to extract covariates for statistical modeling. Duplicated CpG sites on the EPIC array were excluded at random. Eventually, 817 247 CpG sites were included in statistical analyses.

#### Mixed Linear Model on Individual CpG Sites

Using the R package lme4^29^, we calculated mixed linear models on the M-values^30^, with group (psilocybin vs. placebo) as a between and time as a within factor. We included a random effect for patient ID to account for inter-subject variability, one cell type PC and two control probe PCs as covariates, as well as sex, age, smoking, and daily alcohol intake before withdrawal (in units per day). For variance decomposition^31^ and multicollinearity analysis, see Supplementary Figures 1 and 2, respectively. We report genome-wide significance at a suggestive threshold of *p* = 1e-5, as done in previous research^32^.

Primary effects of interest in the linear model were the interaction effects time2*group and time3*group, indicating significant differences in change from baseline methylation between treatment and placebo groups, 24 h and 28 days after the intervention, respectively. Significant longitudinal effects were *post hoc* tested by running cross-sectional *t*-tests for the relevant time points (T2 or T3, respectively) on the beta values adjusted for the covariates (using *R*’s predict function). We report goodness-of-fit for models with significant effects as variance explained by all regressors in the model^33^ (conditional R^2^ calculated with performance library^34^). Model estimates resulting in singular fits were excluded from subsequent analyses, leaving 649 975 CpG sites in the dataset. Results were annotated using the manufacturer’s manifest (https://emea.support.illumina.com/array/array_kits/infinium-methylationepic-beadchip-kit/downloads.html). Non-annotated CpGs that appeared as relevant in the *post hoc* tests, *i.e.,* significant effects with a magnitude of |Δβ| > 0.02 for the cross-sectional difference, were also screened on https://ewascatalog.org/ for associated genes.

#### Sensitivity Analysis

A sensitivity analysis using G*Power 3.1^35^ to estimate the effect size required to detect a deviation of total explained variance R^2^ in a linear multiple regression model was calculated. Parameters used here were α = 1e-5, *n* = 40, Power = 0.8, and number of predictors = 9.

#### Downstream Analyses

Differentially methylated regions (DMR) were detected using the dmrff algorithm^36^ (maximal gap size: 1 000 bp; *p*-cutoff: 0.05). Visualization was based on the qqman package^37^ and used the *p*-values for longitudinal effects. We considered DMRs with a minimum coverage of two CpG sites and report the results of cross-sectional *post hoc* tests for the affected CpGs.

Weighted Correlation Network Analysis (WGCNA)^38^ was performed on the 5% most variable CpG sites (45 962 CpGs) from the two post-intervention timepoints to derive co-methylation modules that capture potential treatment effects. Networks were constructed using the following parameters: soft power threshold = 3 (defined by the criterion of approximate scale-free topology: truncated R^2^ > 0.90), minimum module size = 50, mergeCutHeight = 0.25, and maxBlockSize = 46 000. In WGCNA, modules are labeled by colors. The module’s eigen-CpGs (analogous to eigengenes^38^), representing a weighted average of the module’s expression profile, were calculated and correlated with phenotypic variables of interest: group, duration of abstinence, and mean alcohol intake during the 4-week post-treatment period, as well as Δ-values (T3-T1) for BDI and BHS. Eigen-CpG methylation values were winsorized to two standard deviations. For each variable, we report the module with the strongest correlation, including the relation between module membership (correlation between methylation of CpG site and module’s eigen-CpG) and CpG significance (-log(*p*) of correlation between CpG methylation and trait of interest)^38^.

Furthermore, we performed Gene Ontology Overrepresentation Analysis (GO ORA) using missMethyl^39^. This was done on the CpG sites included in the co-methylation modules we report.

#### Candidate Gene Analysis

We also examined methylation changes at CpG sites in a selection of candidate genes chosen based on their proposed involvement in AUD and/or the effects of psychedelics. This selection comprised (i) receptors presumably involved in the effects of or targeted by psilocin, the active metabolite of psilocybin^40^, namely *HTR2A, HTR1A, SLC6A4, NTRK2, GRM2, DRD1,* and *DRD2*^41–46^; (ii) immediate early genes (IEGs) and plasticity-related genes associated with addictive disorders and/or psychedelic drug action: *EGR1, EGR2, FOSB, JUND, BDNF* and *SV2A*^46–54^; and a more heterogeneous group (iii) consisting of genes related to inflammation (*TNF, IL6, CXCL8),* glutamate (*GRIN2B*) and glucocorticoid (*NR3C1*) signaling, as well as epigenetic regulation (*HDAC2*) that are implied to play a role in AUD^55–58^.

330 CpG sites annotated to candidate genes were retrieved and screened for nominally significant longitudinal effects (*p* < 0.05). Significant CpGs were *post hoc* tested for cross-sectional differences at T2 or T3, respectively, using *t*-tests on β-values adjusted for the covariates from the linear model. We report CpGs with significant (*p* < 0.05) cross-sectional differences that exceeded |Δβ| = 0.02.

Furthermore, we screened for baseline differences in the candidate CpGs between treatment responders (<1 standard unit of alcohol during 4-week follow-up; *n* = 6) and non-responders (≥1 standard unit of alcohol during follow-up; *n* = 11). Due to the small sample sizes for this comparison, non-parametric Wilcoxon tests were chosen, and results for *p* < 0.05 uncorrected are reported.

#### Mediation Analysis

We conducted a mediation analysis to explore whether changes in methylation mediated the effects of psilocybin treatment on depression scores (ΔBDI/ΔBHS), focusing on eight CpG sites with prior significance. Methylation changes (ΔT2-T1 or ΔT3-T1) were modeled by group (mediator model), and ΔBDI/ΔBHS was modeled by methylation and group (outcome model). Methylation values were adjusted using prior mixed model predictions. As methylation showed no significant effect in the outcome models, mediation analysis was not pursued further.

## Results

### Clinical Outcomes

A detailed description of the clinical results can be found in Rieser *et al.*, 2025^21^. Drinking-related outcome metrics did not show significant differences: duration of abstinence had a mean of 11.24 days after placebo versus 17.77 days after psilocybin (*p* = 0.095); mean daily alcohol intake was 1.39 units after placebo and 0.84 units after psilocybin (*p* = 0.331). We also analyzed group differences in ΔBDI and ΔBHS (Δ: T3-T1) and found significant effects: ΔBDI = −0.41 in the placebo group vs. ΔBDI = −6.18 after psilocybin, as well as ΔBHS = 0.65 in the placebo group and ΔBHS = −1.59 after psilocybin. These differences represent large effect sizes reaching statistical significance with *p* = 0.017 and Cohen’s d = 0.87 for ΔBDI and *p* = 0.017 and Cohen’s d = 0.86 for ΔBHS.

### EWAS Results

Linear modeling identified one intergenic CpG site with a significant interaction effect (*p* < 1e-5; Figure 2a & Sup. Tab. 1.1) between group and time point 2; cg04492946 (*p* = 1e-6). However, *post hoc* testing revealed no significant cross-sectional difference (*p* = 0.714). At T3, 17 CpG sites showed significant longitudinal effects (*p* < 1e-5; Figure 2b & Sup. Tab. 1.2). *Post hoc* analysis of cross-sectional group differences identified four CpGs with an effect size of |Δβ| > 0.02 (negative Δβ indicates lower methylation in psilocybin group): cg01405499 (*p* = 0.017; Δβ = −0.02; R^2^ = 0.74), cg23107740 (*p* = 1.1e-7; Δβ = −0.04; R^2^ = 0.35), cg09767929 (*p* = 9.9e-11; Δβ = 0.03; R^2^ = 0.38), and cg17174681 (*p* = 2.1e-12; Δβ = 0.02; R^2^ = 0.43). None of these CpG sites were annotated to genes in the Illumina manifest. Using https://ewascatalog.org/, however, cg23107740 was annotated to transducin-like enhancer of split 4 (*TLE4)* and cg01405499 to the non-coding RNA *LINC01250.* Of note, cg23107740 displayed a significant group difference at baseline, which was inverted at time point 3 (Sup. Fig. 3).

**Figure 2:**
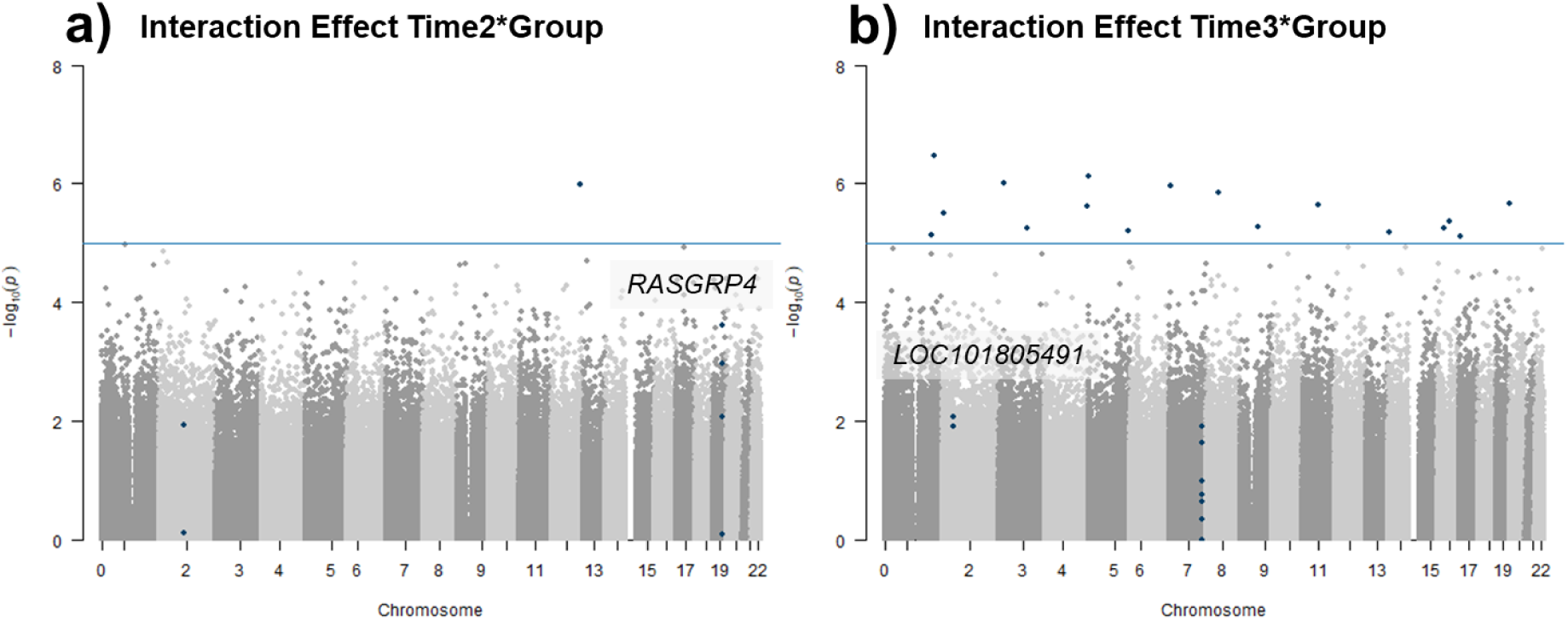
Results of linear modeling and DMR analysis. **a)** Manhattan plot showing the −log(*p*)-values for the longitudinal effects time2*group at each CpG against its location in the genome. Blue line indicates genome-wide significance cutoff of *p* < 1e-5. Blue dots indicate CpGs that reach genome-wide significance or belong to a DMR. Where possible, DMRs are annotated to genes. **b)** Same as a) but for longitudinal effects time3*group.

### Sensitivity Analysis

Sensitivity analysis revealed a minimal effect size of *f^2^* = 1.54 for our models. Based on *R*^2^ = *f*^2^/(1 + *f*^2^) a model fit needs to explain a proportion of at least R^2^ = 0.8 in the methylation values to produce accurate findings with a power of 0.8 in our study.

### Differentially Methylated Regions (DMR)

Two DMRs showed psilocybin-dependent longitudinal effects at T2 and T3, respectively. DMRs are highlighted in Figures 3a and 3b. One DMR associated with psilocybin-dependent changes at T2 was intergenic (*n* = 2 CpGs; z = 5.49; *p_FDR_* = 0.026), the other one covered CpGs in the gene RAS (rat sarcoma) guanyl nucleotide-releasing protein 4 (*RASGRP4*) (*n* = 4 CpGs; z = 6.23; *p_FDR_* = 3.2e-4). Longitudinal psilocybin-dependent effects at T3 also covered two regions: one intergenic DMR (*n* = 7 CpGs; z = −6.47; *p_FDR_* = 6.6e-5) and one in the non-coding RNA *LOC101805491* (*n* = 2 CpGs; z = −6.08; *p_FDR_* = 8e-4; annotated using https://ewascatalog.org/). Cross-sectional *post hoc* tests of the covered CpGs revealed that only one affected methylation site displayed a significant cross-sectional effect of |Δβ| > 0.02. This was cg14565721 in *RASGRP4* gene showing hypermethylation 24 h after psilocybin (*p* = 0.041; Δβ = 0.02; R^2^ = 0.88). The four CpG sites in this DMR lie within a 2 000 bp distance from the transcription start site of *RASGRP4*, suggesting potential involvement in transcription regulation.

**Figure 3:**
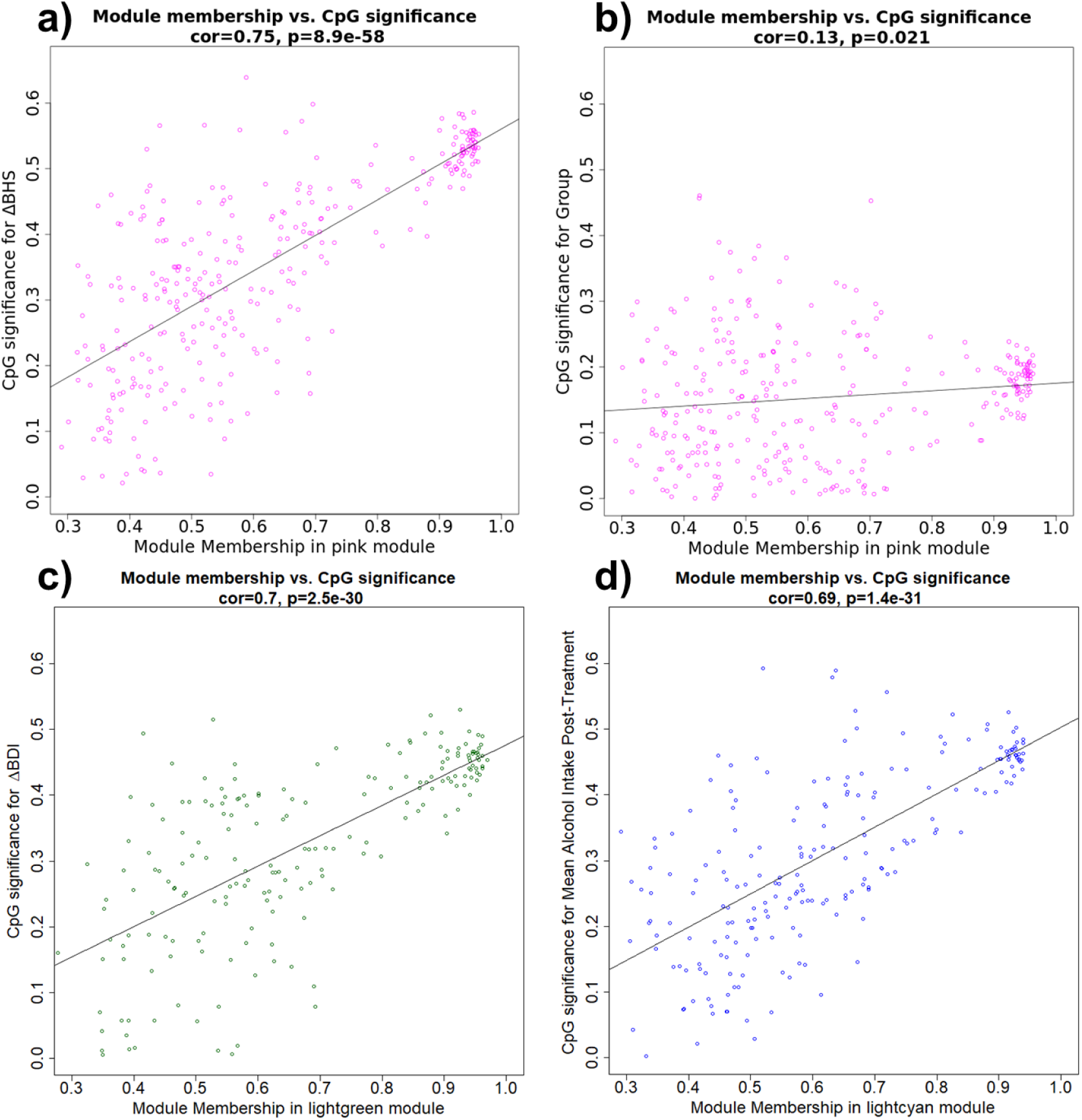
Co-methylation modules with significant correlation between CpG significance for variable of interest and module membership. Colors indicate the co-methylation module. The regression line indicates the strength of the association. **a)** Membership pink module vs. CpG significance ΔBHS. **b)** Membership pink module vs. CpG significance group. **c)** Membership lightgreen module vs. CpG significance ΔBDI. **d)** Membership lightcyan module vs. CpG significance mean alcohol intake during 4-week follow-up.

### Weighted Correlation Network Analysis (WGCNA)

WGCNA identified 34 co-methylation modules (median size: *n* = 206; range: *n* = 73-17612). Significant correlations between module eigengenes and the variables of interest occurred in the following modules: pink (group: *r* = −0.24, *p* = 0.044; ΔBHS: *r* = 0.56, *p* = 5.2e-7), lightgreen (ΔBDI: *r* = 0.48, *p* = 2e-5), lightcyan (mean alcohol use: *r* = 0.47, *p* = 3.5e-5), green (duration of abstinence: *r* = 0.27, *p* = 0.021). For most modules, the relationship between modules and the respective variables was confirmed by strong correlations between module membership and gene significance for the CpG sites within the modules^38^ (Fig. 3): pink (group: *r* = 0.13, *p* = 0.021; ΔBHS: *r* = 0.75, *p* = 8.9e-58), lightgreen (ΔBDI: *r* = 0.7, *p* = 2.5e-30), lightcyan (mean alcohol use: *r* = 0.69, *p* = 1.4e-31). Only for the green module was this correlation insignificant (duration of abstinence: *r* = 0.06, *p* = 0.2).

### Gene Ontology Overrepresentation Analysis (GO ORA)

GO term analyses on the co-methylation modules revealed enrichment of terms broadly related to neurodevelopment and (lightgreen module, Sup. Tab. 2.1), immune function and cell cycle regulation (pink module, Sup. Tab. 2.2), synaptic transmission and intracellular protein regulation (lightcyan module, Sup. Tab. 2.3), as well as calcium signaling and gene/protein regulation (green module, Sup. Tab. 2.4), among other functions. However, no enrichment survived correction for false discovery rate (FDR).

### Candidate Analysis

Among the 330 target CpG sites investigated, 19 reached nominal significance (*p* < 0.05) for the interaction effect of time and group at T2 and 16 for the interaction effect at T3 (see Sup. Tab. 3.1 & 3.2). None of these effects remained significant after FDR correction. Among the significant CpGs, three sites showed significant cross-sectional effects exceeding |Δβ| = 0.02 (see Fig. 4): cg01620540 (T2: *p* = 4.8e-4, Δβ = −0.03; T3: *p* = 1.6e-3, Δβ = −0.03, R^2^ = 0.56) and cg27068143 (T2: *p* = 0.049, Δβ = −0.03, R^2^ = 0.75), both of which lie within 2 000 bp up/downstream of the transcription start site of *HTR2A,* as well as cg11484872 (T2: *p* = 7e-3, Δβ = −0.03, R^2^ = 0.54) which is associated with the *TNF* promoter. Again, promoter association/proximity to the transcription start site of the CpG sites implies potential regulatory functions.

**Figure 4:**
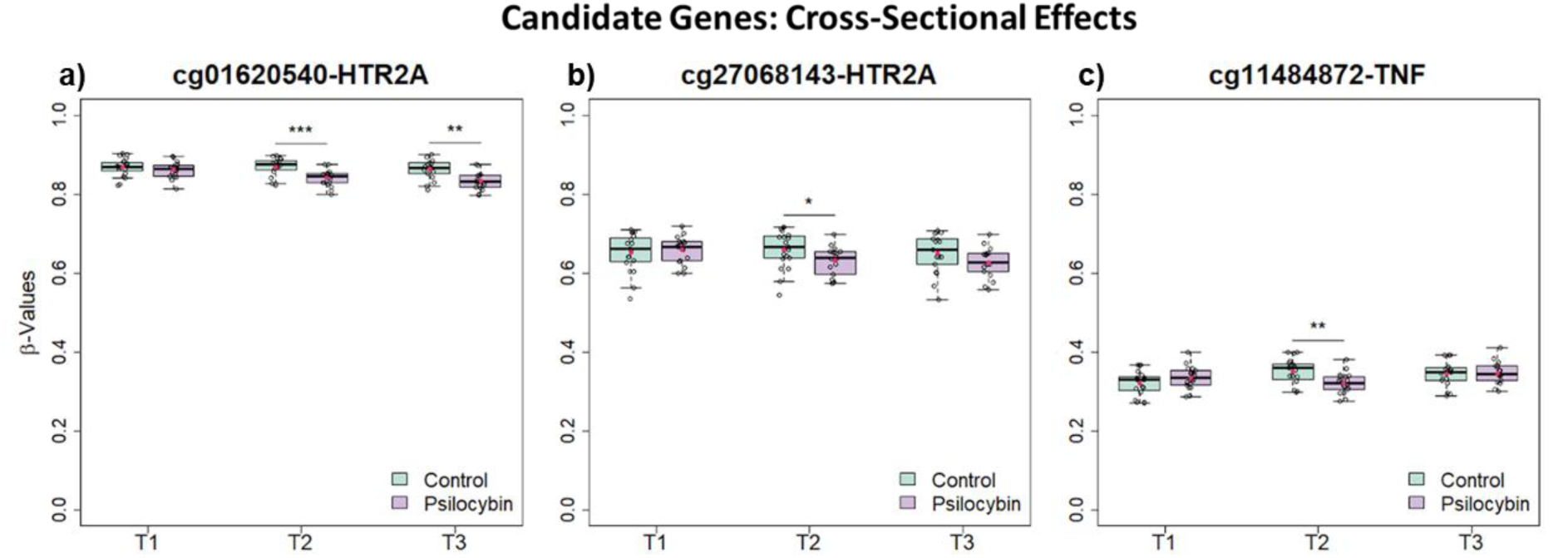
CpG sites that showed significant cross-sectional effects in *post-hoc* testing of candidate genes. Green is the placebo group, violet is the psilocybin group. *: *p* < 0.05; **: *p* < 0.01; ***: *p* < 0.001. **a)** cg01620540 in the *HTR2A* promoter. **b)** cg27068143 in the *HTR2A* promoter. **c)** cg11484872 in the *TNF* promoter.

Next, we compared methylation levels of the candidate CpGs between responders (abstinent until 4-week follow-up) and non-responders in the psilocybin group. We identified 12 CpGs that displayed nominally significant differences before the psilocybin treatment (Tab. 1).

**Table 1:** Significant (*p* < 0.05) baseline differences between treatment responders and non-responders. *p*-values stem from Wilcoxon-tests. Δβ**-**values represent the contrast responders > non-responders. FDR column shows multiple comparison corrected *p*-values. Gene annotation derived from the manufacturer’s manifest. Location describes the position of the CpG sites within the gene: TSS = (distance from) transcription start site; i.e., TSS1500 describes a CpG site within 1

| CpG | *p* | $\Delta\beta$ | FDR | Gene | EntrezID | Location |
| --- | --- | --- | --- | --- | --- | --- |
| cg12067298 | 0,037 | -0,028 | 0,988 | BDNF | 627 | TSS1500 |
| cg15710245 | 0,007 | 0,030 | 0,800 | BDNF | 627 | TSS1500 |
| cg11501905 | 0,027 | 0,069 | 0,988 | DRD1 | 1812 | exon |
| cg19730798 | 0,048 | -0,005 | 0,988 | DRD2 | 1813 | TSS200 |
| cg00105415 | 0,015 | 0,076 | 0,800 | EGR2 | 1959 | TSS1500 |
| cg02881684 | 0,048 | -0,072 | 0,988 | FOSB | 2354 | exon |
| cg24609211 | 0,027 | -0,052 | 0,988 | FOSB | 2354 | exon |
| cg15287403 | 0,015 | 0,007 | 0,800 | GRM2 | 2912 | exon |
| cg00221070 | 0,037 | -0,014 | 0,988 | HTR1A | 3350 | exon |
| cg08652028 | 0,010 | 0,049 | 0,800 | NR3C1 | 2908 | exon |
| cg13848734 | 0,007 | 0,010 | 0,800 | NTRK2 | 4915 | TSS1500 |
| cg14312898 | 0,015 | 0,011 | 0,800 | SLC6A4 | 6532 | TSS1500 |

### Mediation Analysis

Upon testing the eight statistically and biologically relevant CpG sites identified in our analyses on psilocybin treatment, we observed no significant effects for methylation in the outcome models for either ΔBDI or ΔBHS, rendering mediation analysis obsolete.

## Discussion

We present the first methylome-wide exploration of psilocybin-induced changes in blood DNA methylation in a clinical population (*N* = 40). In this analysis, we identified a number of CpG sites and co-methylation modules with potential relevance for psilocybin’s effects that may support future hypothesis-driven research.

In our EWAS, four CpG sites showed significant methylation changes after psilocybin. One of these was annotated to a gene, *TLE4*, where it is located in the gene body^59^. *TLE4* is a transcriptional co-regulator involved in developmental^60–62^ and immunoregulatory^63,64^ processes. Furthermore, *TLE4* regulates maturation and maintenance of corticothalamic projection neuron identity^62^, Schwann cell differentiation^65^, and post-synaptic gene transcription at neuromuscular junctions^66^. *TLE4* has also been implicated in addictive behavior in a preclinical study on oxycodone self-administration^67^. Given this context and the suggested role of structural plasticity in psilocybin’s therapeutic effects^68^, there may be a relationship between psilocybin-induced alterations in *TLE4* methylation and potential neuroplastic effects of psilocybin in AUD.

Furthermore, we discovered four DMRs associated with psilocybin-induced methylation changes. One DMR implicated in effects at T2 covered a gene, *RASGRP4.* This signaling molecule contributes to the development of mast cells^69^ as well as the regulation of immune responses^70,71^. Psychedelics, including psilocybin, possess immunomodulatory capacities^72^ and reductions in neuroinflammation may contribute to their lasting psychological benefits^73^. In AUD, on the other hand, (neuro)inflammatory processes are upregulated^74^, seemingly exacerbating cognitive symptoms associated with this condition^75^. Methylation changes in *RASGRP4* may reflect, at least in part, psilocybin’s immunomodulatory effects.

WGCNA revealed several co-methylation modules associated with either treatment group or the drinking-/depression-related psychometrics. The pink module showed significant correlation with the treatment, as well as with one of the psychometric measures (ΔBDI), and included loci relevant to immune function and cell cycle regulation. This, again, indicates a relationship between the potential effects of psilocybin on the immune system and its anti-depressive capacities, as suggested by previous research^73^. The other modules correlating with ΔBHS, duration of abstinence, and mean alcohol use during the 4-week follow-up did not relate to the psilocybin treatment. It is known that DNA methylation patterns in AUD change during prolonged abstinence^76,77^, for instance, involving gene loci related to immune function^78^ and neuroplasticity^79^. As the modules covered biological processes related to synaptic transmission and gene transcription, they possibly describe such abstinence-related methylation changes that are independent of psilocybin treatment.

The candidate gene analysis revealed evidence for hypomethylation in two CpG sites within the promoter of *HTR2A*, which codes for the primary molecular target of psychedelics, the 5HT2a receptor. Aberrant methylation of *HTR2A* has been associated with psychiatric symptoms such as impulsivity in cocaine use disorder^80^, or depressive rumination in people suffering from adverse childhood experiences^81,82^. Normalization of such *HTR2A* methylation might thus lead to symptom relief in patients suffering from conditions like depression or AUD. We also observed a transient hypomethylation in a CpG site in the *TNF* promoter. This may represent a temporary influence on immune signaling. Interestingly, psilocybin decreases *TNF* blood levels in the short term^73^.

Lastly, our investigation of baseline differences between responders to psilocybin treatment (abstinent at 4-week follow-up) and non-responders revealed nominally significant differences in several CpG sites related to neuronal plasticity (*BDNF*, *NTRK2*, *EGR2*, *FOSB*) and various neurotransmitter systems (*DRD1*, *DRD2*, *GRM1*, *HTR1A*, *SLC6A4*, *NR3C1*). The search for predictors of psychedelic treatment responsivity is ongoing and currently focuses on phenomena like the acute effects of psilocybin on brain activity and phenomenology, or changes in language patterns shortly after substance intervention^83,84^. Less is known about molecular factors that could predict treatment responses before psychedelics are administered, and the genes identified here might provide a starting point for future research focusing on this question.

This study is subject to some limitations. One being the small sample size. Thus, it is not too surprising that the data we present here did not survive multiple comparison corrections, and most model fits remain below our R^2^ threshold for sufficient power. Furthermore, on average, the effect sizes we observed are small. Consequently, it appears that a single administration of psilocybin did not cause strong and persistent effects on DNA methylation.

Despite these limitations, we present potential novel epigenetic associations with psilocybin treatment for AUD patients, indicators for methylation changes in genes involved in serotonin and immune signaling, and possible methylomic predictors of treatment responsivity. Further research is necessary to corroborate the findings.

## Acknowledgements

We thank Fabian Streit for providing us with the processing pipeline for the genotype data.

## Author Contributions

MU planned the details of the analysis, modified and executed most of the code for the data processing, and prepared the manuscript. LZ supervised the planning and execution of the analysis, contributed and revised code for the preprocessing and the analysis, and assisted in the preparation of the manuscript. NR planned and executed the clinical study, including withdrawal of the blood samples and acquisition of the psychometric data, prepared the psychometric data for this analysis, and assisted in the preparation of the manuscript. MH and KP supervised the planning and execution of the clinical study and contributed to the planning of the analysis. FV supervised the planning and execution of the clinical study. RS and MWM planned the epigenetic study and contributed to the planning of the analysis and the editing of the manuscript. All authors have read and agreed upon the final version of the manuscript.

## Data Availability

Data will be made available on the European Genome Archive (EGA).

## Supplementary Material

**Sup. Fig. 1:**
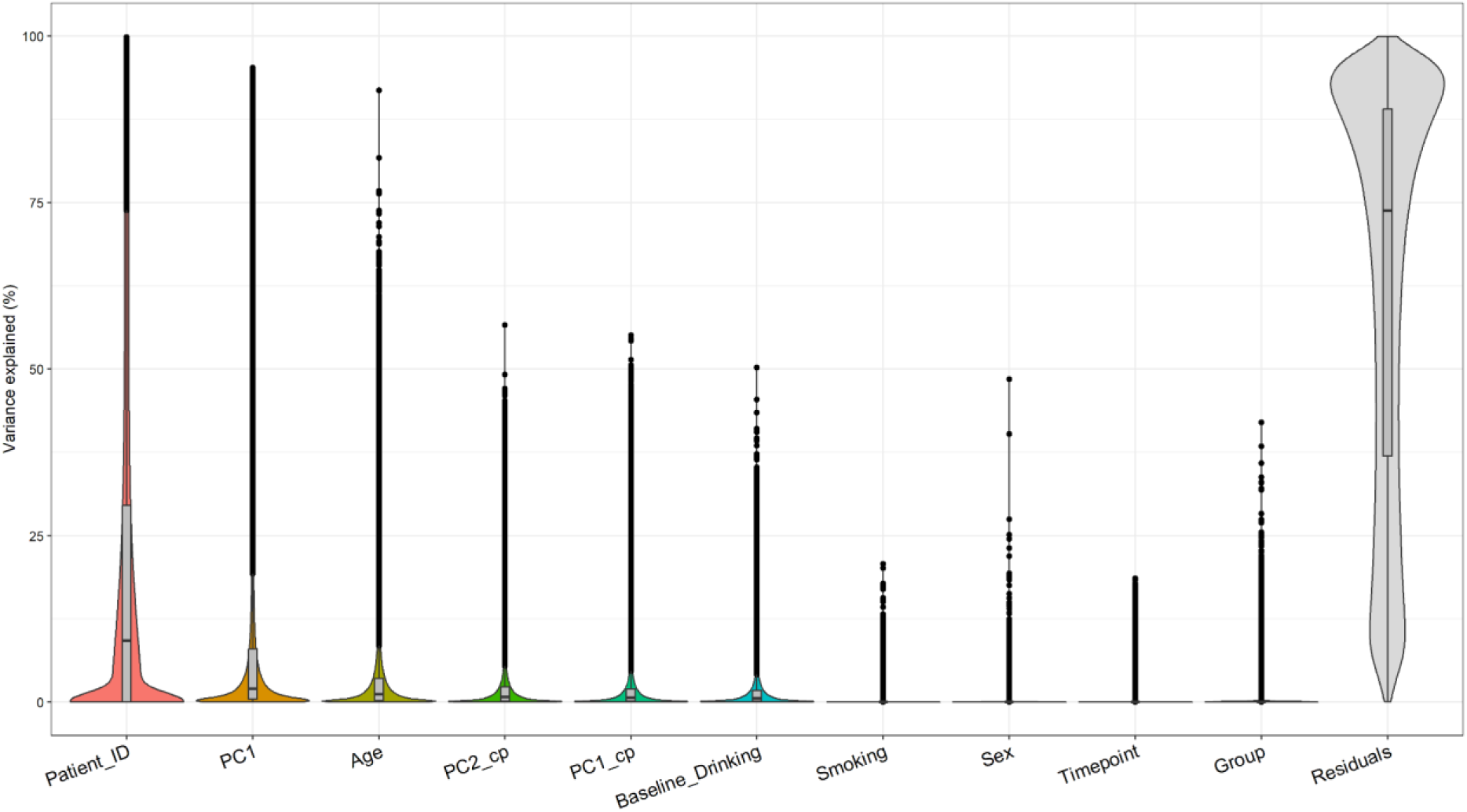
Variance decomposition with covariates included in linear model. PC1: cell type principal component 1; PC1/2_cp: control probe principal component 1/2.

**Sup. Fig. 2:**
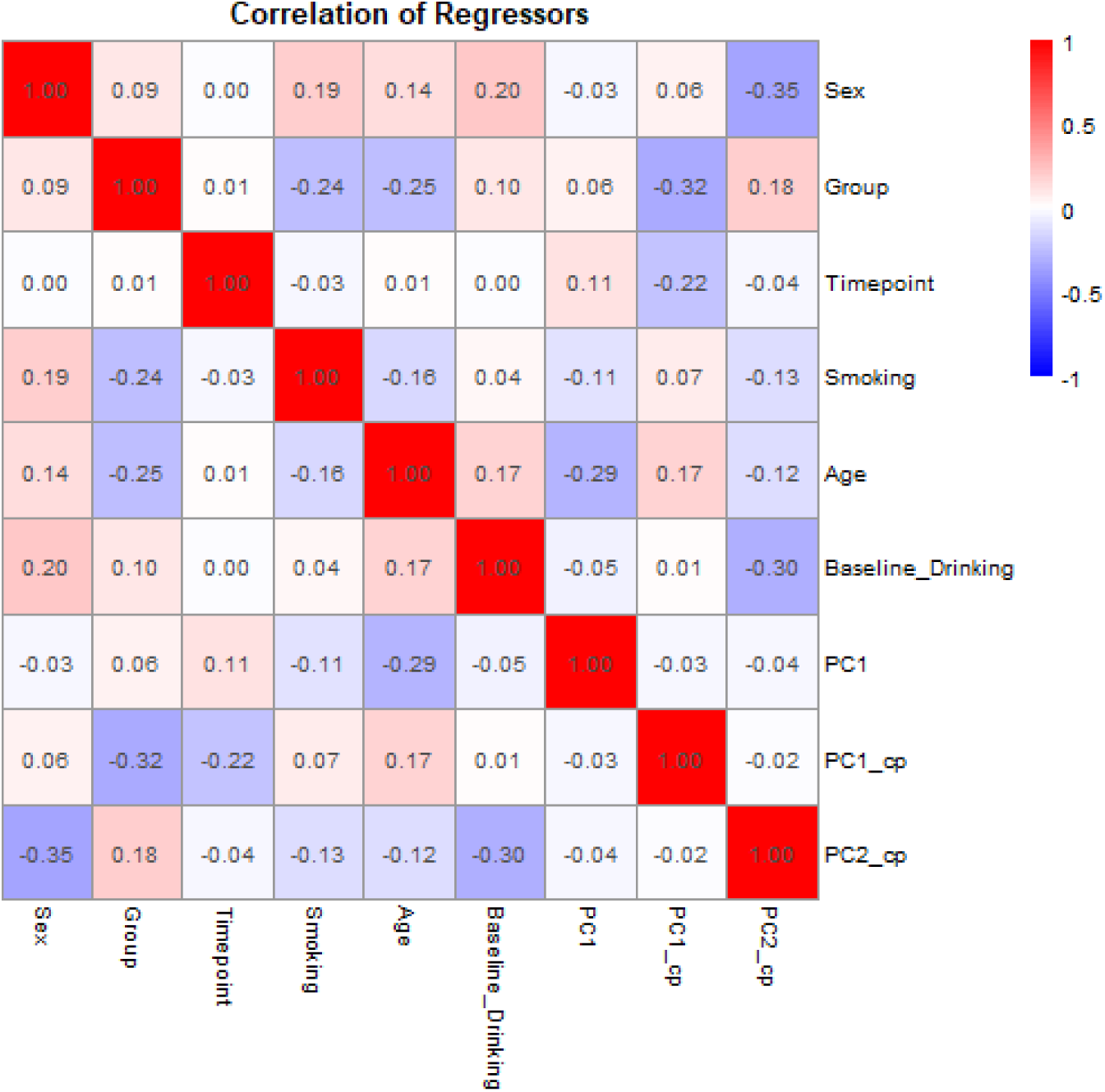
Pearson correlation matrix with regressors included in the model. PC1: cell type principal component 1; PC1/2_cp: control probe principal component 1/2.

**Sup. Fig. 3:**
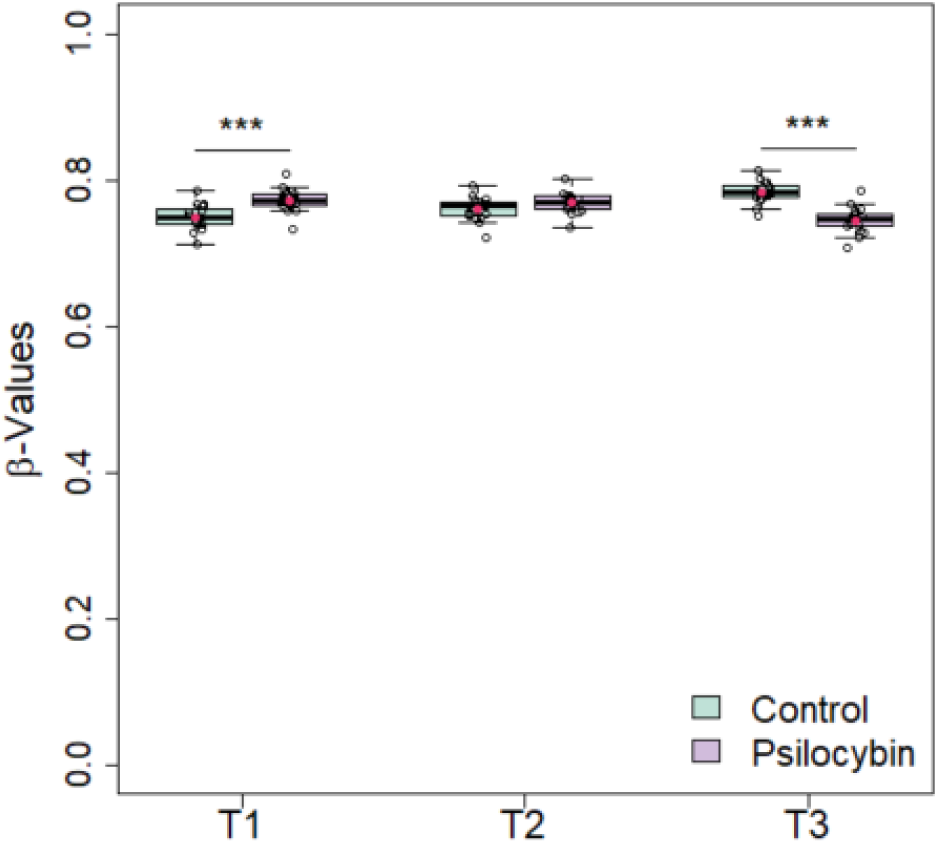
Cross-sectional effects of cg23107740 methylation. Already at baseline, there was a random group difference that inverted throughout the experimental procedure.

**Sup. Fig. 4:**
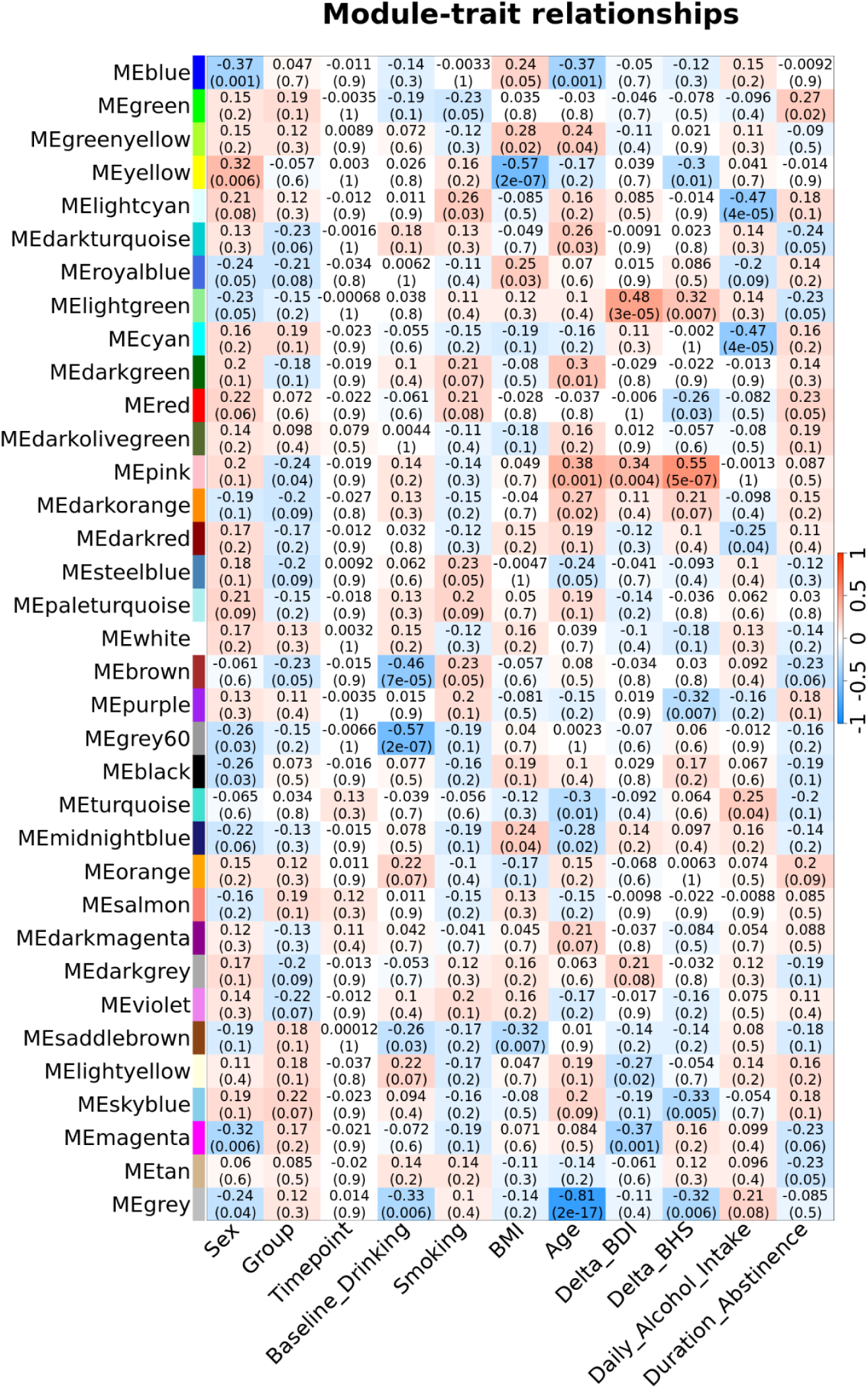
Pearson correlation matrix for com-methylation module eigen-CpGs and phenotypic traits. Upper value in each cell is Pearson’s *r*, lower value is *p*. Delta_BDI: change in Beck’s Depression Inventory between T1 and T3; Delta_BHS: change in Beck Hopelessness Scale between T1 and T3.

